# Linking plant-soil feedback with resource competition to understand plant coexistence

**DOI:** 10.64898/2026.09.13.751226

**Authors:** Shengman Lyu, Xiangyu Liu, Jake M. Alexander

**Author notes:** Correspondence: Department of Ecology and Evolution, University of Lausanne, 1015 Lausanne, Switzerland. **Author contributions:** S.L. and J.A. conceived the study and designed and performed the field and greenhouse experiments. S.L. performed the data analyses and wrote the first draft of the manuscript. J.A. and X.L contributed to interpreting the results, provided revisions, and approved the final version of the manuscript. **Code and data availability**: All data and R code used to reproduce the analyses and figures in this study are publicly available on GitHub: https://github.com/ShengmanLyu/Plant-soil-feedback-and-plant-coexistence.

## Abstract

1. Plant–soil feedback (PSF) is increasingly recognized as an important mechanism influencing plant coexistence, but most studies have examined PSF in isolation under controlled conditions. Under natural conditions, PSF operates together with other processes such as resource competition, yet how these processes jointly shape plant coexistence across environmental gradients remains poorly understood.
2. We combined field and greenhouse experiments to assess the contribution of PSF to plant coexistence at low- and high-elevation sites in the Swiss Alps. In the field, we quantified total niche and fitness differences and predicted interaction outcomes for nine plant species pairs, integrating PSF, resource competition, and other processes. In a complementary greenhouse experiment, we used soil microbial communities conditioned during the field experiment to isolate PSF-driven niche and fitness differences for the same species pairs.
3. PSF alone generated strong fitness differences but weak niche differences in the greenhouse experiment, predominantly predicting competitive exclusion. Interaction outcomes predicted from PSF alone corresponded poorly with those in the field, suggesting that PSF in isolation poorly explain plant coexistence under natural conditions. Nevertheless, PSF-driven niche and fitness differences were related to those quantified in the field, which were likely driven largely by resource competition, suggesting that PSF may contribute to coexistence by reinforcing or counteracting other ecological processes. These relationships varied between elevations, highlighting the context dependence of PSF contributions to coexistence.
4. *Synthesis.* Our study shows that the impacts of plant–soil feedback on plant coexistence cannot be understood from their effects in isolation. Instead, their contribution depends on how they interact with other ecological processes and on the environmental context in which these processes operate. Together, our findings emphasize the need to move beyond studying PSF in isolation towards quantifying how it combines with other mechanisms that drives coexistence under natural conditions.

## INTRODUCTION

Understanding the processes that drive species coexistence and maintain community diversity is a central goal in community ecology (Hutchinson 1961; Tilman 1988; Chesson 2000). Traditionally, studies have focused primarily on interactions between plants and their abiotic environments, such as competition for limiting resources (Tilman 1988; Hautier *et al*. 2009). Recently, interactions between plants and soil microbes, such as plant-soil feedback (PSF), are increasingly recognized as an additional mechanism shaping plant coexistence (Bever *et al*. 1997; Bever 2003; Mordecai 2011; Kandlikar *et al*. 2019; Ke & Wan 2019; Chung *et al*. 2023a; Xi *et al*. 2025). PSF occurs when a plant alters soil communities, which in turn influence the population growth of itself and its neighbouring species (Bever *et al*. 1997; van der Putten *et al*. 2013). Although the effects of PSF on plant coexistence are supported by many empirical studies, most have tested PSF under controlled greenhouse conditions and in isolation from other processes (reviewed by Petermann *et al*. 2008; Crawford *et al*. 2019; Yan *et al*. 2022). However, in natural communities, PSF operates together with other processes such as resource competition, and so far, how these processes jointly shape plant coexistence under natural remains poorly understood. Addressing this knowledge gap may help explain the exceptionally high plant diversity maintained across steep environmental gradients in mountain ecosystems (Körner 2021).

Coexistence theory predicts that the outcomes of plant interactions depend on the balance between niche differences, which promote stable coexistence, and fitness differences, which favour competitive exclusion (Chesson 2000). Both resource competition and plant– soil feedback (PSF) can contribute to niche and fitness differences through distinct mechanisms (Illustrated in the Box 1) (Kandlikar *et al*. 2019; Ke & Wan 2020). In natural communities, they may jointly influence plant coexistence in at least two ways. First, PSF may act independently of resource competition and other processes, such that their effects on niche and fitness differences combine additively to determine interaction outcomes. In this case, their relative contributions depend on the strength of PSF relative to other processes and may vary across environments. For example, resource competition may be weaker under colder or less productive conditions (Callaway *et al*. 2002; Hargreaves 2024), where soil microbes may have stronger effects on plant performance due to slower nutrient cycling and greater reliance on microbially mediated resource acquisition (Florianová C Münzbergová 2025). PSF may also contribute relatively more under such conditions, such that PSF-driven interaction outcomes will more closely resemble those observed in the field. Conversely, under warmer and more productive conditions where resource competition dominates, PSF may contribute less and PSF-driven outcomes may correspond poorly with field interaction outcomes (Lekberg *et al*. 2018). Yet, it remains unclear how well PSF measured in isolation explain plant coexistence under natural conditions, and whether their contribution varies across contrasting environments.

In addition to independent contributions, PSF may interact non-additively with resource competition and other processes by modifying the niche and fitness differences they generate, thereby reinforcing or counteracting their effects on coexistence (Fig. 1c, d). First, PSF may modify niche differences between competitors. For example, when species with different resource niche dimensions are also associated with distinct host-specific pathogens because of their different evolutionary histories or functional strategies (Xi *et al*. 2021; Sweeney *et al*. 2025), PSF reinforces niche differences arising from resource partitioning and could strengthen stabilization and shift interacting species from competitive exclusion towards coexistence (green arrow in Fig. 1d). Conversely, when species with different resource niche share general soil mutualists that reduce their biotic niche differentiation (Xi *et al*. 2025), PSF could counteract resource-driven niche differences, thereby weakening stabilization and potentially shifting coexisting species towards competitive exclusion (purple arrow in Fig. 1d).

**Figure 1.**
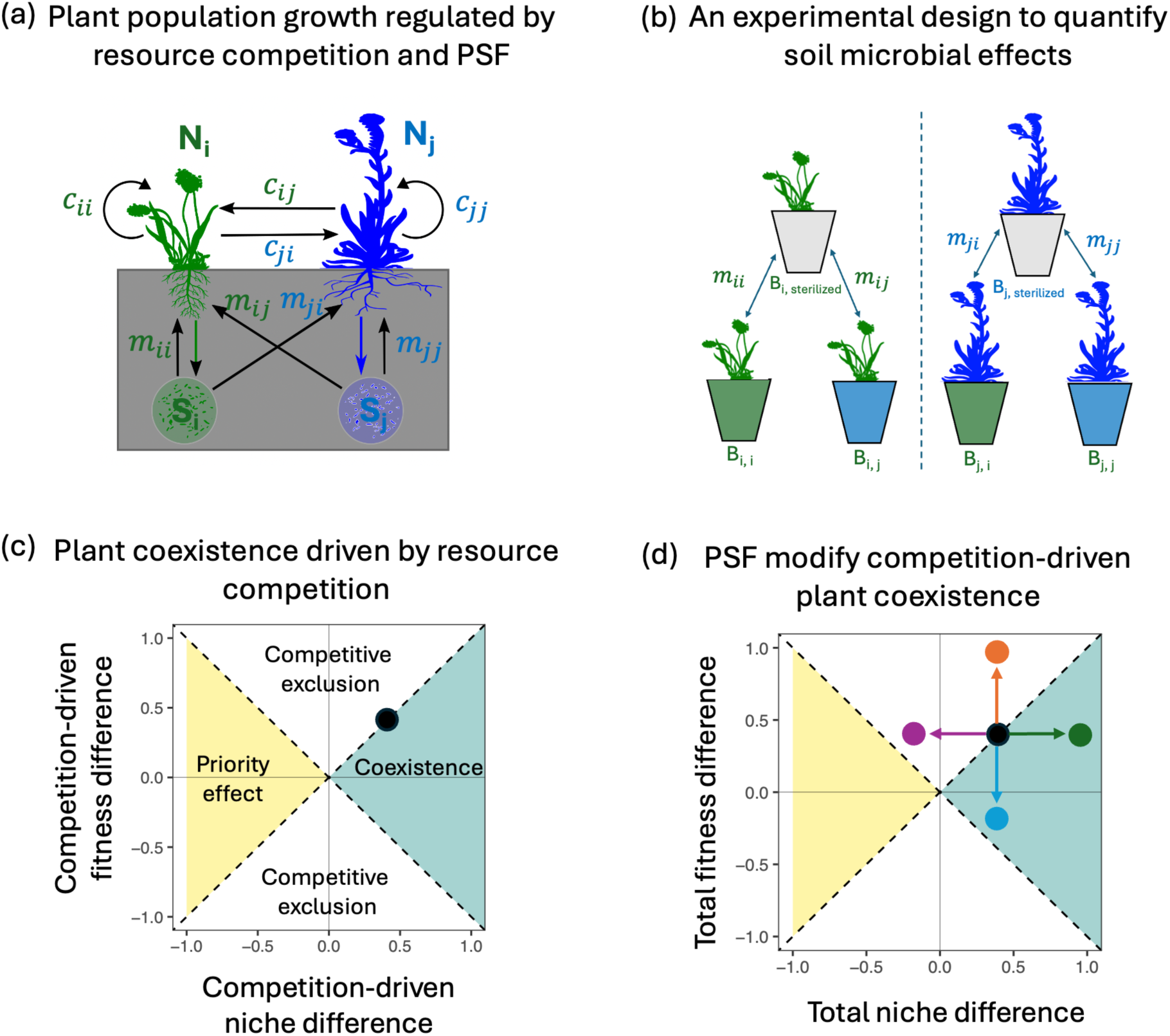
Conceptual framework linking plant–soil feedback (PSF) and resource competition to plant coexistence. (a) Population growth of two interacting plant species regulated by resource competition (represented by *c*) and PSF (represented by *m*). (b) Experimental design used to quantify soil microbial effects on plant growth. *B_i, j_* represents the biomass of species *i* grown with soil microbes conditioned by species *j*, *B_i, i_* represents the biomass of species *i* grown with soil microbes conditioned by itself, *B_j,i_* represents the biomass of species *j* grown with soil microbes conditioned by species *i*, and *B_j, j_* represents the biomass of species *i* grown with soil microbes conditioned by itself. *m_i, j_, m_i, i_, m_j, i_* and *m_j, j_* represents the microbial effect, quantified by comparing biomass in conditioned versus sterilized soil. (c) Interaction outcomes driven by resource competition are determined by the resulting niche and fitness differences. The black dot represents a species pair at the boundary between stable coexistence and competitive exclusion. (d) PSF can modify the niche and fitness differences generated by resource competition, shifting species pairs from their initial position (black dot) towards different interaction outcomes. Arrows illustrate possible PSF-induced changes in total niche and fitness differences, with their lengths indicating the strength of PSF relative to other processes.

Similarly, PSF may also generate fitness differences that reinforce or counteract those arising from resource competition and other processes. For example, although PSF alone often generates fitness differences that favour competitive exclusion (Box 1) (Kandlikar *et al*. 2020; Yan *et al*. 2022), their consequences depend on which competitor benefits. If soil microbes disproportionately benefit the weaker resource competitor, for example through a trade-off between resource competition ability and dependence on or responsiveness to soil mutualists (Romero *et al*. 2023), PSF could reduce fitness differences and shift species towards coexistence (blue arrow in Fig. 1d). Conversely, if the stronger resource competitor also benefits more from soil microbes (Yang *et al*. 2026), PSF could amplify fitness differences and intensify competitive exclusion (orange arrow in Fig. 1d). Consequently, whether PSF promotes stable coexistence or competitive exclusion cannot be solely understood from PSF-driven niche and fitness differences in isolation. Yet PSF effects on coexistence have primarily been studied in isolation (Petermann *et al*. 2008; Yan *et al*. 2022), and we know little about how PSF-driven niche and fitness differences relate to those arising from resource competition and other processes.

In this study, we combined a field competition experiment with a complementary greenhouse PSF experiment to assess the contribution of PSF to plant coexistence under natural conditions and how this contribution varies across an elevational gradient. In the field, we parameterized integral projection models (IPMs) to quantify field-derived total niche and fitness differences and predicted interaction outcomes for 18 plant species pairs at low- and high-elevation sites, integrating the effects of PSF, resource competition, and other processes operating in the field. In the greenhouse, we used soils conditioned during the field experiment to isolate PSF effects and quantify PSF-driven niche and fitness differences for the same species pairs under climatic conditions approximating those in the field. By comparing the two experiments, we assessed both how well PSF alone captures interaction outcomes in the field and how PSF-driven niche and fitness differences relate to those emerging from all processes operating under natural conditions.

We addressed three questions: (1) How well do interaction outcomes predicted from PSF alone correspond with those observed in the field? (2) How are PSF-driven niche and fitness differences related to the total niche and fitness differences observed in the field? (3) Do the correspondences and relationships vary between low- and high-elevation environments? Resource competition is often a major determinant of plant coexistence (Tilman 1988; Hautier *et al*. 2009), and previous work showed that competition for light and soil resources strongly predicted competitive ability and coexistence among the study species, particularly at low elevation (Lyu & Alexander 2024). We therefore expected PSF-driven outcomes to correspond poorly with field outcomes at low elevation, where resource competition is likely to dominate, but more closely at high elevation, where resource competition is weaker and PSF may contribute relatively more. Following the same reasoning, we expected PSF-driven niche and fitness differences to correspond more closely with total niche and fitness differences measured in the field at high than at low elevation.

## MATERIALS AND METHODS

### Field competition experiment

A detailed description on the field experiment and population modelling can be found in (Lyu & Alexander 2022). Briefly, the field experiment was established in two sites along an elevation gradient in the western Swiss Alps in autumn 2017 (Table S1). The low and high sites were located at 1400 and 1900 m above the sea level, with mean annual temperatures of 5.9 and 2.5 °C, respectively (Table S1 in the Supplementary Information). We selected six dominant perennial species in the study area, including two legumes: *Medicago lupulina* (Melu) and *Anthyllis vulneraria ssp. alpestris* (Anal), and four forbs: *Plantago lanceolata* (Plla), *Plantago alpina* (Plal), *Aster alpinus* (Asal), and *Salvia pratensis* (Sapr) (Table S2). Seeds were obtained from regional commercial suppliers (Table S2). We selected nine species pairs among all the 30 (6 x 5 species) possible pairwise combinations spanning a wide range of functional trait differences.

We first established high-density monocultures for each species in each site in spring 2017, then transplanted focal individuals raised in a greenhouse for four weeks into the established monocultures in autumn 2017. The focal individuals were planted 10 cm apart to reduce competition among focal individuals. We also transplanted focal individuals to non-competition plots (bare ground) to measure plant performance in the absence of competition (20 cm apart). We followed all individuals between 2017 and 2020 to estimate individual vital rates, including annual survival, growth, flowering and seed production. In 2019, we performed a separate experiment in each site to estimate seed germination and seedling establishment.

We used integral projection models to incorporate individual vital rates into estimates of population growth rates (λ). We estimated intrinsic population growth in the absence of competition (λ_intrinsic_) using individuals grown in the non-competition plots. We estimated invasion population growth (λ_invasion_) when a focal species is at low density (i.e. without intraspecific competition) and its competitor is at its single-species equilibrium density using individuals grown in the high-density monocultures of competitors.

### Coexistence in the field

We quantified niche and fitness difference and predicted competitive outcomes for each species pair using estimates of λ_intrinsic_ and λ_invasion_ from the field experiment following Carroll et al. (Carroll *et al*. 2011), which is theoretically equivalent to Chesson’s decomposition approach (Chesson 2000)(Box 1). Because these field estimates integrate PSF, resource competition, and other processes operating under field conditions, we refer to them as total niche and fitness differences (Fig. 1d). For a pair of species *i* and *j*, we first calculated the sensitivity of species *i* competing against species *j* as:

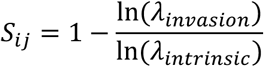

where λ_invasion_ is population growth rate of species *i* invading the monoculture of its competitor j at equilibrium density, and λ_intrinsic_ is the intrinsic growth rate species *i* in the absence of competition. Sensitivity is positive for competitive interactions, and greater sensitivities indicate stronger competition. A pair of species with smaller niche differences (i.e. greater niche overlap) experiences stronger interspecific competition, therefore larger mean sensitivities (Chesson 2000; Godoy & Levine 2014). Therefore, niche overlap, ρ, can be calculated as the geometric mean of the two species’ sensitivities and their niche difference then equals one minus niche overlap:

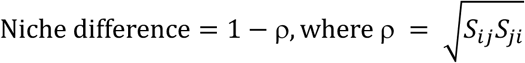

Fitness differences quantify the degree of asymmetry in species’ competitive abilities (Chesson 2000; Godoy & Levine 2014). A pair of species with large fitness differences experience different intensities of interspecific competition, which can be calculated as the geometric standard deviation of sensitivities:

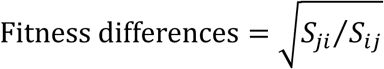

ND can be rearranged as -log(p) and FD as log(FD) on a natural logarithm scale (e.g.(Willing *et al*. 2024)). A superior competitor is predicted to exclude the inferior competitor if their fitness differences are sufficient to overcome the stabilizing effects of niche differences, that is, when FD > ND. Otherwise, when FD < ND, fitness differences are insufficient to drive competitive exclusion, and only stable coexistence or priority effects are possible. When ND > 0, species have stabilizing niche differences and stable coexistence occurs; when ND < 0, species have destabilizing niche overlap and priority effects occur, meaning that whichever species is initially established within a community has an advantage and excludes the other.

### Greenhouse PSF experiment

We established a greenhouse experiment in 2020 to isolate the effects of soil microbial communities on plant growth and quantify PSF-driven niche and fitness differences for the same species pairs examined in the field competition experiment. In total, the experiment consisted of 372 plants: (6 sterilized + (6 intraspecific + 22 interspecific combinations) x 2 sites) x 6 replicates. We grew each of the six species in sterilized field soil either without living soil microbes or inoculated with soil microbial communities conditioned during the field experiment (Fig. 1b). For each focal species, we used soil conditioned by itself and the species with which it was interacted in the field experiment, resulting in 6 intraspecific and 22 interspecific species–soil combinations. We separately used microbial communities conditioned at the low- and high-elevation sites to assess whether microbial effects varied with elevation. Each species–soil combination was replicated six times.

The background potting soil was collected near the low-elevation field site, thoroughly mixed, sieved through a 1-cm mesh to remove stones and living plant material, and autoclaved twice at 120 °C, with >24 h between cycles. Living soil inoculum was collected at the end of the 2020 growing season from high-density monocultures of each species at both field sites. Soil was collected where no heterospecific plants occurred within a 10-cm radius, sieved through a 0.5-cm mesh to remove stones and living plant and animal material, and thoroughly mixed within each monoculture treatment. Inocula were stored at 4 °C and used within 48 h of collection.

Plants were grown individually in pots (8 cm diameter × 10 cm depth), each containing approximately 500g of soil. Sterilized-control pots contained 90% sterilized background soil and 10% (by mass) sterilized field soil comprising equal proportions of soil from all monoculture treatments. Live-inoculum pots contained 90% sterilized background soil and 10% living soil from the corresponding monoculture treatment. Seeds were surface-sterilized in 10% bleach for 10 min, rinsed with deionized water, and germinated on filter paper moistened with deionized water in Petri dishes. Seedlings were transplanted individually into pots as soon as radicles emerged.

Greenhouse temperatures were programmed to approximate average growing-season conditions across the two field sites, with a 12 h day/12 h night cycle and temperatures of 20 °C during the day and 10 °C at night. Pots were arranged in six blocks, watered with deionized water three times per week, and repositioned weekly to minimize spatial environmental heterogeneity. After three months, plants were harvested at ground level, oven-dried at 60 °C for one week, and weighed to determine dry aboveground biomass. We used dry aboveground biomass as a proxy for population growth because plant size is strongly associated with survival and reproduction in our study species (Lyu & Alexander 2022), and individual growth contributes more (ca. 67%) to population growth under competition than other vital rates (Lyu & Alexander 2023).

### PSF-driven coexistence in the greenhouse experiment

We computed PSF-driven niche and fitness differences using metrics derived by Bever and colleagues (Bever *et al*. 1997; Bever 2003) and Kandlikar and colleagues (Kandlikar *et al*. 2019; Kandlikar *et al*. 2021), as illustrated in the Box 1. For a pair of species *i* and *j*, we first quantified the effect of soil microbes on plant growth relative to sterilized soil using log-transformed aboveground biomass of plants grown in the greenhouse experiment:

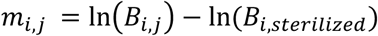

Where *B_i_*_,j_ and *B_i_*_,*sterilized*_ are the aboveground biomass of plant *i* grown in soil microbes conditioned by plant j and sterilized soil, respectively (Fig. 1). Positive *m* values mean that plant growth is enhanced by soil microbes (e.g. by mutualists), while negative values mean that growth is reduced by soil microbes (e.g. by pathogens). We then calculated PSF-driven niche and fitness differences following the Equations 5 and 6, respectively (Box 1). Depending on the relative magnitude of PSF-driven niche and fitness differences, PSF can result in three possible outcomes: stable coexistence, competitive exclusion and priority effects, for which the conditions are same as those driven by resource competition as described above.

### Statistical analyses

#### Effects of soil microbes on plant growth

We first tested how the effects of soil microbial communities on plant growth varied among focal and conditioning species and between conditioning elevations. We used linear mixed-effects models with aboveground biomass as the response variable and included focal species, conditioning species, elevation, and all two- and three-way interactions as fixed effects, and block as a random effect. We used a similar model structure to test how intraspecific and interspecific PSF (*m*) varied among the six species and between elevations. In both cases, we simplified the model by sequentially removing non-significant interaction terms while retaining all main effects, and assessed the significance of fixed effects and retained interactions in the final models.

#### Linking PSF-driven with field-derived coexistence

We tested agreement in competitive outcomes quantified between the field and greenhouse PSF experiments using χ^2^ contingency table tests, treating interaction outcomes as categorical responses (i.e. whether a pair of species can coexist or not). Analyses were performed separately for the two study sites to allow for site-specific environmental effects. We tested whether PSF-driven niche and fitness differences were correlated with field-derived total niche and fitness differences using linear mixed-effects models. We first included site as a covariable and, in case the relationships were significantly different in the two sites, we ran the model separately for each site. Because each species interacted with several species, we included the identify of both the focal and competitor species as random factors to account for this nonindependence. We used the lme4 package to fit all linear mixed-effects models (Bates *et al*. 2015) and used type-II *F*-tests to test the significance of fixed effects using the *car* package (Fox & Weisberg 2011). All analyses were performed in R 4.6.1 (R Core Team 2026).

## RESULTS

### Effects of soil microbes on plant growth

Overall, plants grew better in soil inoculated with living microbial communities than in sterilized soil, although negative microbial effects occurred in a few species–soil combinations (Fig. 2). Specifically, *Salvia pratensis* (Sapr) showed negative responses to microbial communities conditioned by *Medicago lupulina* (Melu) and *Plantago lanceolata* (Plla), while *Plantago alpina* (Plal) showed a negative response to its conspecific-conditioned microbial community at the high-elevation site (Fig. 2). Microbial effects on plant growth differed strongly among focal species (*F_5, 300_* = 444.01, *P* < 0.001; Fig. 2). *Aster alpinus* (Asal) and *P. lanceolata* (Plla) benefited most from soil microbes, producing, on average, more than four times as much biomass in inoculated as in sterilized soil. *M. lupulina* (Melu) and *Anthyllis alpestris* (Anal) produced more than twice as much biomass, whereas microbial effects were weaker for *P. alpina* (Plal) and *S. pratensis* (Sapr) (Fig. 2).

**Figure 2.**
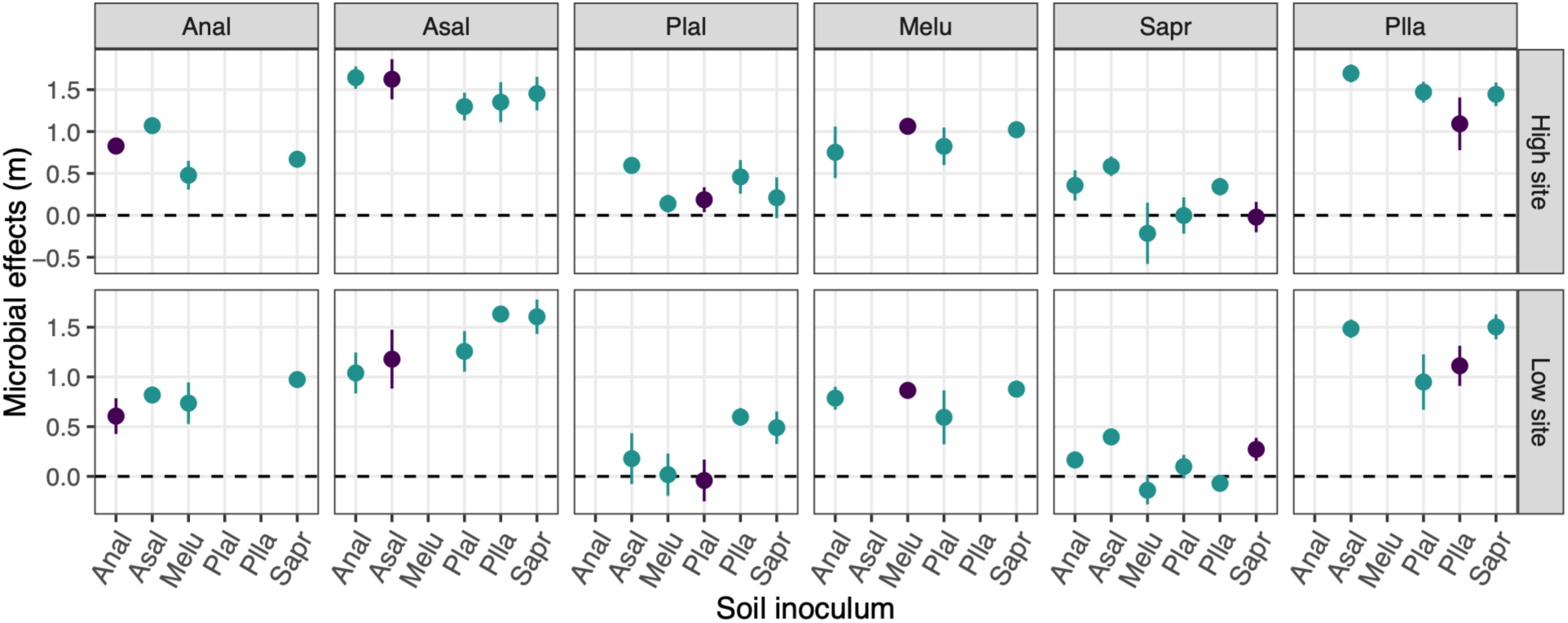
Effects of soil microbial communities on plant biomass, quantified as the log response ratio of biomass in inoculated versus sterilized soil (microbial effect, *m*). Each panel represents a focal species, with green and black points indicating interspecific (*m_ii_*) and intraspecific (*m_i_*_j_) microbial effects, respectively. Positive values indicate greater plant biomass in inoculated than sterilized soil, whereas negative values indicate lower biomass in inoculated soil. Missing points indicate focal–conditioning species combinations that were not included in the experiment. Points show means ± SE.

Microbial effects also differed among conditioning species (*F_5, 300_* = 26.46, *P* < 0.001; Fig. 2). On average, microbial communities conditioned by *A. alpinus* (Asal) had the strongest positive effects on plant growth, followed by those conditioned by *S. pratensis* (Sapr), *P. lanceolata* (Plla), *A. alpestris (Anal)*, *P. alpina* (Plal), and *M. lupulina* (Melu) (Fig. 2). However, there was no significant interaction between focal and conditioning species (*F_17, 300_* = 20.98, *P* = 0.2227), indicating little evidence that the effects of microbial communities depended on the identity of the focal species. Consistent with this weak host specificity, microbial effects did not differ between intraspecific and interspecific soil communities (*F_1, 300_* = 2.49, *P* = 0.115; Fig. 2). Finally, microbial communities conditioned at high elevation had, on average, slightly more positive effects on plant growth than those conditioned at low elevation (*F_1, 300_* = 4.54, *P* = 0.033), and this elevational difference varied among conditioning species (conditioning species × elevation: *F_5, 300_* = 15.46, *P* = 0.009).

### PSF-driven coexistence versus coexistence in the field

When considering PSF alone, competitive exclusion was predicted for 17 of the 18 species pairs across the two elevations (Fig. 3a,b). PSF generated relatively strong fitness differences but weak niche differences, such that niche differences were generally insufficient to overcome fitness differences. The only exception was the *P. alpina–S. pratensis* pair at low elevation, for which PSF-driven niche differences exceeded the corresponding fitness difference, predicting stable coexistence (Fig. 3a).

**Figure 3.**
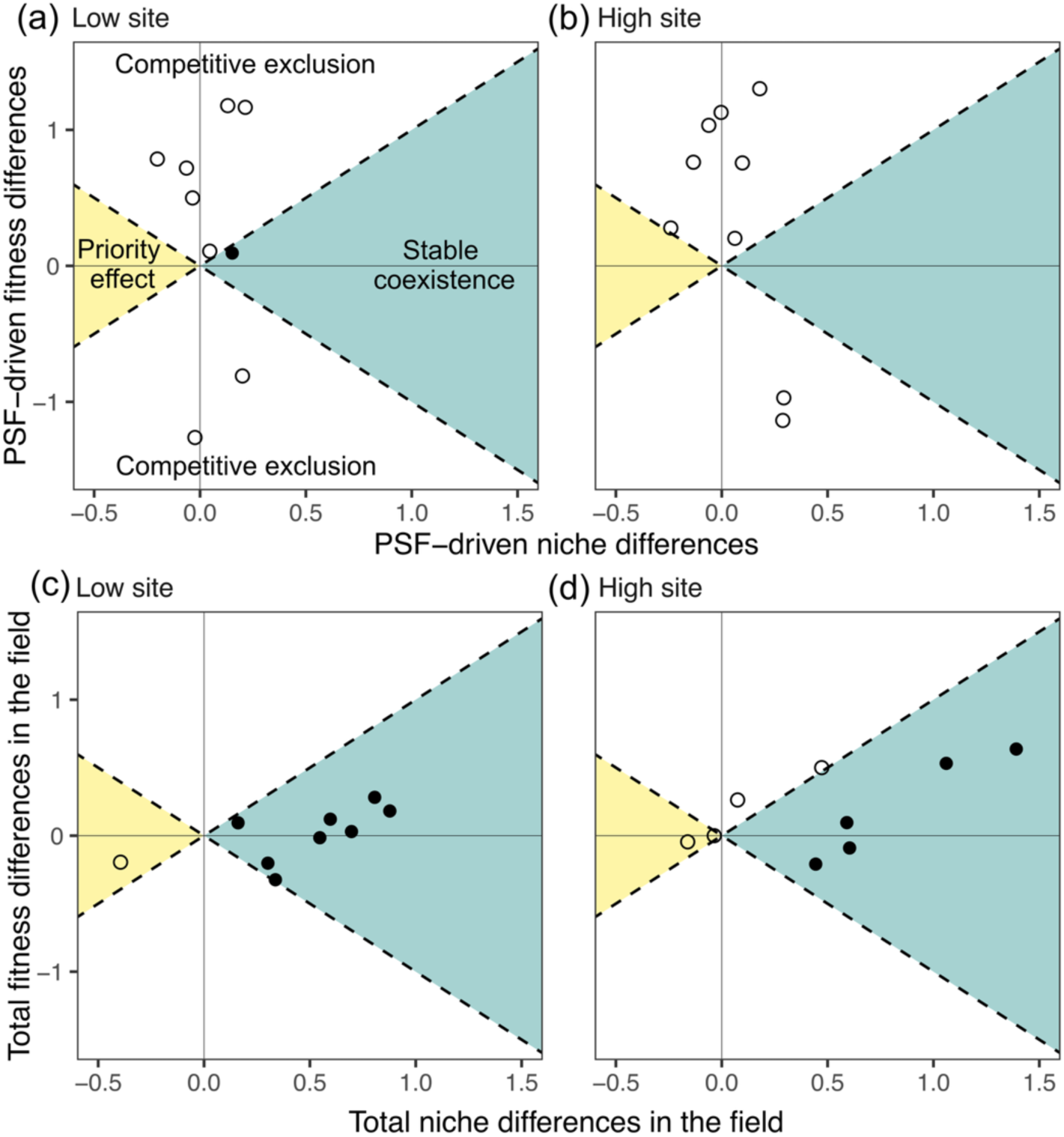
Niche and fitness differences and predicted interaction outcomes based on plant– soil feedbacks (PSF) alone in the greenhouse experiment using soil microbial communities conditioned at the (a) low- and (b) high-elevation sites, and field-derived total niche and fitness differences and interaction outcomes at the (c) low- and (d) high-elevation sites. Each point represents a species pair. Filled points indicate stable coexistence (blue-shaded areas), whereas open points indicate competitive exclusion (unshaded areas) or priority effects (yellow-shaded areas). Dashed lines delimit the regions corresponding to the three interaction outcomes.

In contrast, stable coexistence was predicted for 13 of the 18 species pairs in the field, where the estimated niche and fitness differences integrate PSF, resource competition, and other processes operating under natural conditions (Fig. 3c,d). Field-derived total niche differences were generally more pronounced than PSF-driven niche differences. However, because PSF-driven and field-derived niche and fitness differences were estimated using different experimental approaches and are not directly comparable in absolute magnitude, we did not interpret differences in their numerical magnitudes. Instead, we compared the interaction outcomes they predicted and their relationships across species pairs. Predicted outcomes based on PSF alone differed significantly from field outcomes at low elevation (*χ^2^* test: *χ^2^* = 8, *P* = 0.004), whereas the difference was only marginally significant at high elevation (*χ^2^* = 2.89, *P* = 0.08).

### Relationships between PSF-driven and field-derived total niche and fitness differences

Relationships between PSF-driven niche and fitness differences and their field-derived total counterparts differed between elevations (Fig. 4). At low elevation, PSF-driven niche differences were negatively related to field-derived total niche differences (*F_1,_* _9_ = 34.934, *P* < 0.001; Fig. 4a): species pairs with larger total niche differences in the field tended to have smaller PSF-driven niche differences. The relationship was also negative at high elevation but was not significant (*F_1, S_* = 0.012, *P* = 0.91; Fig. 4b).

**Figure 4.**
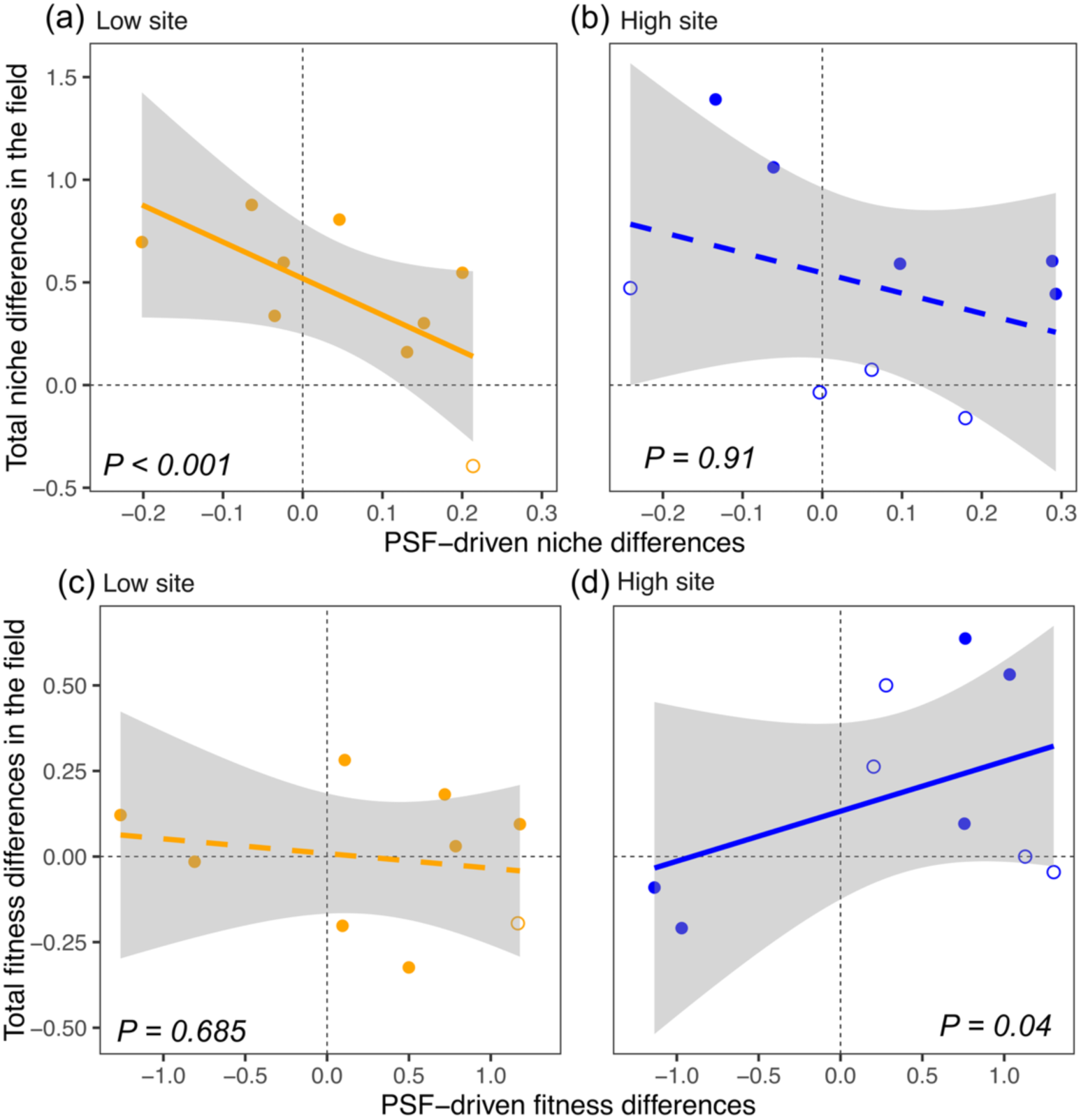
Relationships between PSF-driven niche (a, b) and fitness (c, d) differences and field-derived total niche and fitness differences at the low- and high-elevation sites. Each point represents a species pair, with filled points indicating pairs predicted to stably coexist and open points indicating pairs predicted to undergo competitive exclusion or priority effects in the field experiment (Fig. 3c,d). Lines show fitted linear relationships, with solid lines indicating significant relationships (*P* < 0.05) and dashed lines indicating non-significant relationships (*P* > 0.05). Grey shading indicates 95% confidence intervals around the fitted relationships.

In contrast, PSF-driven fitness differences were unrelated to field-derived total fitness differences at low elevation (*F_1, S_* = 0.165, *P* = 0.685; Fig. 4c), but were positively related at high elevation (*F_1, S_* = 4.205, *P* = 0.04; Fig. 4d). Thus, species pairs with larger PSF-driven fitness differences also tended to have larger total fitness differences in the high-elevation field community, whereas no such relationship was evident at low elevation.

## DISCUSSION

Plant–soil feedback (PSF) has often been invoked as an important driver of plant coexistence (Bever *et al*. 1997; Bever 2003; Mordecai 2011; Kandlikar *et al*. 2019; Ke & Wan 2019; Chung *et al*. 2023a; Xi *et al*. 2025), yet their contribution relative to other processes under natural conditions remains poorly understood. By combining field estimates of coexistence with PSF effects isolated in a complementary greenhouse experiment, we found that interaction outcomes predicted from PSF alone corresponded poorly with those observed in the field. Nevertheless, PSF-driven niche and fitness differences were related to those quantified in the field, but these relationships differed between elevations. In high elevation, PSF-driven niche differences were negative correlated with those quantified in the field, while PSF-driven fitness differences were positive correlated with those quantified in the field in low elevations. Together with the disagreement in interaction outcomes, our results suggest that PSF may play a minor role in driving plant coexistence under the field conditions. Our results also suggest that understanding the role of PSF in natural communities requires considering both environmental context and their relationships with other coexistence processes.

### PSF alone can generate strong fitness differences and often lead to exclusion

Contrary to common expectations, PSF alone was predicted almost exclusively to result in competitive exclusion in the greenhouse experiment, with only a single case of stable coexistence. This was because PSF generated strong fitness differences but only weak niche differences. This pattern closely mirrors the quantitative synthesis of Yan and colleagues, which found across 518 plant species pairs that soil microbes generated substantially stronger fitness differences than stabilizing or destabilizing effects and, consequently, predicted competitive exclusion much more frequently than coexistence (Yan *et al*. 2022). Such large fitness differences imply that one species gains a greater net benefit from soil microbes than its competitor. While such fitness asymmetries could partly reflect a sampling effect arising from the particular species included in our experiment, some species may happen to associate with more beneficial microbial communities than others. Their consistency across studies suggests that they may also reflect inherent differences among species in their responsiveness to soil mutualists or susceptibility to soil pathogens (Xi *et al*. 2021; Yan *et al*. 2022; Romero *et al*. 2023).

In contrast to our expectation, PSF generated only weak niche differences. In the greenhouse experiment, soil microbial communities had almost exclusively positive effects on plant growth, likely reflecting the influence of generalist soil mutualists that enhance resource acquisition (Berendsen *et al*. 2012; Semchenko *et al*. 2022). Again, this pattern is consistent with the study by Yan and colleagues, who found that microbially mediated stabilizing or destabilizing effects were substantially weaker than fitness differences across plant species (Yan *et al*. 2022). Theory predicts that such mutualists generate niche differences only when plant species condition soil microbial communities in ways that benefit their competitors more than themselves (Bever *et al*. 1997; Kandlikar *et al*. 2021), a condition that was rarely met in our study, as microbial effects showed little host specificity. Although stronger niche differences could emerge in more complex environments or over longer timescales (Petermann *et al*. 2008; Crawford *et al*. 2019), the weak host specificity has also been observed in other longer-term field experiments. For example, across two growing seasons in a sagebrush steppe, Chung and colleagues (Chung *et al*. 2023b) found that soil microbes consistently reduced plant growth, but these effects were rarely host-specific, resulting in few negative pairwise PSFs that could stabilize coexistence. Together, our results align with a growing body of empirical work showing that PSF frequently generate strong fitness differences but weak niche differences and more often drive competitive exclusion rather than stable coexistence (Crawford *et al*. 2019; Kandlikar *et al*. 2021; Yan *et al*. 2022).

### PSF may have limited effects on coexistence under field conditions

The outcomes of competition driven solely by PSF in the greenhouse experiment agreed poorly with those quantified in the field, suggesting that PSF might have a limited influence on plant coexistence under natural conditions, at least in our system. This is consistent with recent studies showing that other processes, particularly resource competition, often play a dominant role in shaping plant coexistence in the field and may mask the influence of PSF on plant performance and coexistence outcomes (Lekberg *et al*. 2018; Forero *et al*. 2019; Heinze *et al*. 2020). Consistent with this explanation, a previous study showed that plant traits associated with light competition and soil resource acquisition were strong predictors of population growth under competition and partly explained species niche and fitness differences in the field experiment (Lyu & Alexander 2024).

In addition to the possibility that resource competition and other processes masked or modified PSF effects in the field, our greenhouse experiment may not have fully captured the magnitude or direction of PSF realized under field conditions. First, we inferred PSF effects on coexistence from aboveground growth, whereas soil microbes can influence multiple vital rates, including germination, survival, and reproduction, whose combined effects may have greater effects on population growth (Dostál 2025b). Second, both soil conditioning and feedback effects can accumulate and change over time (Hawkes *et al*. 2013; Ke *et al*. 2021; Liu *et al*. 2025), soils conditioned from the field monocultures may not capture host-specific microbes. Finally, PSF are strongly environment-dependent (Lekberg *et al*. 2018; Dostál 2025a; Florianová & Münzbergová 2025), and controlled greenhouse conditions cannot reproduce the environmental variation and biotic interactions experienced by plants in the field. Indeed, across 36 plant species, Forero and colleagues found no positive relationship between greenhouse- and field-measured PSF, with greenhouse PSF generally stronger in magnitude (Forero *et al*. 2019). Thus, our experiment provides a controlled estimate of how PSF can contribute to niche and fitness differences, rather than a direct estimate of PSF realized in the field. Controlled experiments remain essential for isolating microbial effects, but combining them with direct field measurements will be critical for understanding how PSF contribute to coexistence when multiple ecological processes operate simultaneously (Schittko *et al*. 2016; Lekberg *et al*. 2018; Crawford *et al*. 2019).

### Interactions between PSF- and competition-driven coexistence processes

Although PSF alone poorly explained interaction outcomes in the field, PSF-driven niche and fitness differences were related to those quantified in the field in our experiment. Because field-derived total niche and fitness differences integrate PSF, resource competition, and other processes, these relationships do not allow us to isolate direct interactions between PSF and resource competition. Nevertheless, they suggest that PSF effects are not independent of the other processes generating niche and fitness differences under natural conditions. PSF may therefore contribute to coexistence not only through their effects in isolation, but also by modifying other coexistence processes.

Interestingly, PSF-driven niche differences were negatively related to field-derived total niche differences at both sites, although the relationship was significant only at low elevation (n = 9 species pairs). This negative relationship further suggests that PSF alone did not directly determine the niche differences observed in the field, which would instead be expected to produce a positive relationship. However, PSF generated stronger stabilizing effects among species pairs with weak niche differentiation in the field, suggesting that PSF may stabilize coexistence among species with greater overlap in resource use (Lekberg *et al*. 2018).

In contrast, PSF-driven and field-derived total fitness differences were largely unrelated at low elevation but positively related at high elevation. The lack of a relationship at low elevation suggests that PSF might have had only limited contribution to the total fitness differences observed there. At high elevation, however, the positive relationship suggests that species with greater competitive advantages in the field also tended to benefit more from soil microbes. Such a relationship could arise if competitively dominant species are more effective at recruiting or associating with beneficial soil mutualists (Yang *et al*. 2026).

In this case, PSF could reinforce fitness differences generated by other processes, potentially shifting otherwise coexisting species towards competitive exclusion or further intensifying exclusion. These mechanisms remain hypotheses and require direct measurements of plant resource-acquisition strategies and microbial associations to test (Chung *et al*. 2023a).

More broadly, variation among plant species in resource-acquisition strategies, dependence on soil mutualists, and susceptibility to soil pathogens could determine whether PSF reinforce or counteract niche and fitness differences generated by other processes (Ke & Wan 2020; Xi *et al*. 2025). Together, these findings emphasize that the consequences of PSF cannot necessarily be inferred from PSF-driven niche and fitness differences in isolation, but depend on how they relate to other coexistence processes operating in natural communities.

### Environmental dependence of PSF effects on plant coexistence

The relationships between PSF-driven niche and fitness differences and field-derived total niche and fitness differences differed between sites, suggesting that the contribution of PSF to coexistence is environment-dependent. This may arise from two reasons. First, the relative importance of coexistence processes varies between elevations. Even if PSF effects remain similar, their contribution relative to resource competition may therefore change. For example, a previous work showed that traits associated with resource competition explained a larger amount of the variation in competitive ability at low than high elevation (Lyu & Alexander 2024). Consistent with this pattern, the positive relationship between PSF-driven and field-derived total fitness differences at the high-elevation site suggests that PSF may contribute more to fitness differences where resource competition is weaker. This is consistent with the hypothesis that PSF may be relatively more important in low-productivity environments, such as at high elevations, where resource competition is often weaker (Jiang *et al*. 2024).

Second, PSF effects may themselves vary across environments (Cardinaux *et al*. 2018). In our study, soil microbial communities from the high-elevation site had, on average, more positive effects on plant growth than those from the low-elevation site. This could reflect stronger plant conditioning of beneficial microbial communities or a greater prevalence of mutualistic microbes under relatively harsh conditions such as high-elevation sites (Trivedi *et al*. 2022). Together, these results suggest that environmental conditions may influence both the strength of PSF and their contribution relative to other coexistence processes, leading their effects on plant coexistence to vary across environments (Collings *et al*. 2025).

## CONCLUSION

Our study highlights the potential for plant–soil feedback to influence plant coexistence, primarily by generating strong fitness differences but relatively weak niche differences. This finding calls for a refinement of conventional PSF theory, which has largely emphasized stabilizing effects, and instead highlights fitness differences as an important pathway through which PSF influence coexistence (Kandlikar *et al*. 2019; Yan *et al*. 2022). However, PSF measured in isolation poorly explained interaction outcomes observed in the field, cautioning against directly extrapolating PSF effects from controlled experiments to natural communities. Importantly, PSF-driven niche and fitness differences were related to field-derived total niche and fitness differences, suggesting that PSF may still contribute to coexistence by reinforcing or counteracting other ecological processes (Lekberg *et al*. 2018; Crawford *et al*. 2019). These relationships also varied across environments, highlighting the effects of PSF on coexistence are context dependent (Collings *et al*. 2025). Together, our findings emphasize the need to move beyond studying PSF in isolation towards quantifying how PSF combine with other mechanisms that drive coexistence under natural conditions and across environmental gradients (Heinze *et al*. 2020). Such an approach will be critical for understanding how multiple ecological processes jointly shape plant coexistence, community structure, and biodiversity.

## Supporting information

Table S1 in the Supplementary Information

## Acknowledgements

We thank members of the Plant Ecology Group at ETH Zürich for assistance with field and lab work, in particular Loïc Liberati, Tim Murray, Yan Hess, Jessica Joaquim, Annette Altermatt, and Camille Brioschi. We thank the Commune de Bex for the access to the field sites. S.L. also thanks the Chinese Scholarship Council for financial support (No. 201706100184). J.M.A. received funding from the European Union’s Horizon 2020 research and innovation programme under grant agreement No. 678841.

## Box 1.

### Plant coexistence driven by resource competition and plant-soil feedbacks (PSF)

Here, we use a Lotka–Volterra framework to illustrate how resource competition and plant– soil feedbacks (PSF) jointly influence plant coexistence. The population growth of two interacting plant species *i* and *j*, subject to resource competition (represented by *c*) and PSF (represented by *m*) can be described using a pair of differential equations, one for each species (Fig. 1a):

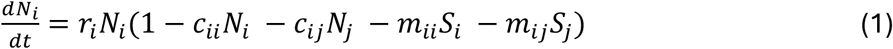

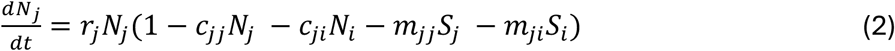

In the absence of competition and PSF, plant populations grow exponentially at their intrinsic growth rates, *r*. We use *c* to represent the per-capita effect of resource competition, with *c_ii_* and *c_i_*_j_ denoting intraspecific and interspecific competition, respectively. We represent soil microbial effects by *m_i_*_j_, the effect on species *i* of soil microbial communities conditioned by species j. Following theoretical PSF models, the abundance of soil microbes associated with a plant species is assumed to increase linearly with the density of that species without a time lag, allowing PSF and resource competition to be represented as plant-density-dependent effects in the same unit (Ke & Wan 2020; Kandlikar *et al*. 2021).

Empirically, *m_i_*_j_ can be estimated by comparing the performance of species *i* with microbial communities conditioned by species j against its performance in a reference soil, such as sterilized soil (Fig. 1b) (Kandlikar *et al*. 2020). This assumes that the measured response reflects population growth, which is reasonable for the study species because plant size is strongly associated with vital rates and growth makes a major contribution to population growth under competition (Lyu & Alexander 2023).

When resource competition acts alone (i.e. setting microbial effects, *m*, to zero), it can influence plant coexistence through two pathways: niche and fitness differences (Chesson & Warner 1981; Chesson 2000). Competition-driven niche differences measure the extent to which species limit themselves more strongly than they limit each other (Equation 3).

Positive niche differences stabilize coexistence and increase as intraspecific competition becomes stronger relative to interspecific competition, for example through resource partitioning (Tilman 1982). Thus, larger competition-driven niche differences make stable coexistence more likely (x-axis in Fig. 1c). Competition-driven fitness differences measure the degree to which one species is competitively superior to the other (Equation 4). Larger fitness differences favour competitive exclusion (y-axis in Fig. 1c) and arise when one species has a greater intrinsic growth rate (the first term in Equation 4) or is less sensitive to competition (the second term in Equation 4) than the other.

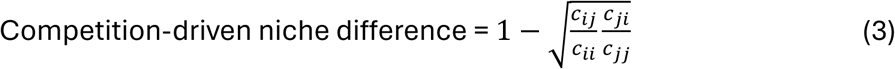

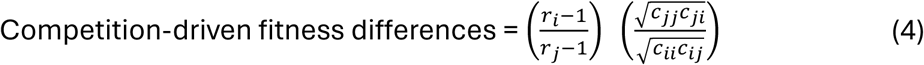

Negative conspecific PSF, for example, through the accumulation of host-specific soil pathogens (e.g. Janzen–Connell effects; (Petermann *et al*. 2008)) can also generate niche differences, whereas PSF that reduce niche differences.

PSF can also influence plant coexistence through the niche and fitness pathways. Following the same reasoning, if the two species on average benefit more (or suffer less in the case of soil pathogens, i.e. negative conspecific PSF) from their competitors’ soil microbial communities than their own (Petermann *et al*. 2008), PSF-driven niche differences are positive and stabilize coexistence (Equation 5). Otherwise, when soil mutualists that preferentially benefit the conditioning plant species (i.e. positive conspecific PSF), PSF-driven niche differences are negative (i.e. increasing niche overlap) and destabilize coexistence. This formulation corresponds to the classic PSF framework developed by Bever and colleagues (Bever *et al*. 1997; Bever 2003). PSF-driven fitness differences measure the degree to which species *i* benefits more (or suffers less) from soil microbes compared to its competitor j (Equation 6). Positive values of fitness differences indicate that species *i* is superior to j, which is expected to be excluded in the absence of niche differences (Kandlikar *et al*. 2019; Kandlikar *et al*. 2021).

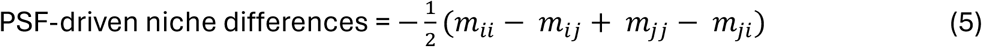

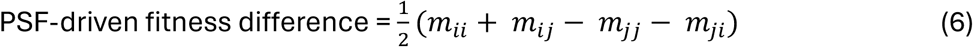

Under natural conditions, resource competition, PSF, and other processes operate simultaneously, combining to generate total niche and fitness differences that determine interaction outcomes (Fig. 1d). Stable coexistence occurs when total niche differences are sufficiently large to overcome total fitness differences; otherwise, one species excludes the other, while negative niche differences can generate priority effects. Consequently, PSF can alter interaction outcomes by reinforcing or counteracting the niche and fitness differences generated by resource competition and other processes (Fig. 1d); the different ways in which PSF may combine with other coexistence processes are discussed further in the Introduction.

**---- End of Box 1 ----**

## Notes

https://github.com/ShengmanLyu/Plant-soil-feedback-and-plant-coexistence

### Competing Interest Statement

The authors have declared no competing interest.

