## Supplementary material for "Linking plant-soil feedback with resource competition to understand plant coexistence": Table S1 in the Supplementary Information

**Table S1**. Environmental characteristics of the two study sites. Long-term mean annual temperature was derived from climate interpolations based on data from 1981-2015. Mean annual temperature of air (15 cm above ground surface) and soil (8 cm below ground surface), and mean soil water content (8 cm below ground surface) were recorded using TMS-4 dataloggers (TOMST®, Czech Republic; https://tomst.com/web/en/) and averaged between July 2019 to July 2020. The number of days covered by snow were calculated as the number of days of which daily mean temperature at the ground surface was between -0.5 and 0.5 °C.

| **Variable** | **Solalex (low site)** | **Anzeindaz (high site)** |
|---|---|---|
| Elevation (m a.s.l.) | 1400 | 1900 |
| Latitude (°N) | 46.2866 | 46.2875 |
| Longitude (°E) | 7.1338 | 7.1687 |
| Long-term mean annual temperature (°C) | 5.9 | 2.5 |
| Mean annual air temperature (°C) | 7.9 | 5.9 |
| Mean annual soil temperature (°C) | 6.5 | 4.9 |
| Mean volumetric soil water content | 0.410 | 0.438 |
| Number of days under snow | 129 | 163 |

**Table S2.** Species included in this study.

| Species | Code | Family | Functional group | Growth form | Elevation origin | Seed supplier |
|---|---|---|---|---|---|---|
| Medicago lupulina | Melu | Fabaceae | Legume | Perennial | lowland | Otto Hauenstein Samen |
| Plantago lanceolata | Plla | Plantaginaceae | Forb | Perennial | lowland | UFA SAMEN |
| Salvia pratensis | Sapr | Lamiaceae | Forb | Perennial | lowland | Otto Hauenstein Samen |
| Anthyllis vulneraria ssp. alpestris | Anal | Fabaceae | Legume | Perennial | highland | Kaertner Saatbau |
| Aster alpinus | Asal | Asteraceae | Forb | Perennial | highland | Jellito |
| Plantago alpina | Plal | Plantaginaceae | Forb | Perennial | highland | Schutz Filisur |
